# Recurrent neural network dynamics may not purely reflect cognitive strategies: A commentary on Ji-An et al. (Nature, 2025)

**DOI:** 10.1101/2025.10.30.685524

**Authors:** Kentaro Katahira

**Affiliations:** Human Informatics and Interaction Research Institute, National Institute of Advanced Industrial Science and Technology (AIST), Tsukuba 305-8566, Japan

## Abstract

One of the fundamental goals of behavioural science is to explain seemingly complex human and animal behaviour in terms of simple mechanistic principles. Ji-An, Benna, & Mattar^1^ (henceforth, JBM) recently proposed a novel analytic and interpretative framework toward this goal. By employing tiny recurrent neural networks (RNNs) with a small number of degrees of freedom, they developed a method for extracting the dynamics of learning and decision-making. They demonstrated that tiny RNNs achieve superior predictive performance compared to classical cognitive models such as reinforcement learning (RL) and Bayesian inference models, and that dynamical systems analyses of the RNNs can reveal cognitive dynamics that differ from those assumed in the cognitive models. This framework is highly promising; however, we suggest allocating room for discussion as to whether the dynamics of RNNs purely reflect the cognitive processes at work in the individual’s brain. In particular, RNNs may adapt to within-individual state changes (e.g., transitions between engaged and disengaged states) in ways not captured by cognitive models, thereby improving model fit without necessarily representing cognitive strategies per se. This raises the possibility that the dynamics of RNNs may include processes that deviate from, rather than directly reflect, individual cognitive strategies.

---

Learning and choice behaviour have traditionally been modelled using cognitive computational models, such as RL and Bayesian inference models^2–4^. However, these models are constrained by their dependence on human-devised hypotheses. More flexible neural network models, particularly recurrent neural networks (RNNs) that can capture temporal dependencies in choice sequences, have attracted increasing attention^5–7^. RNNs often fit behavioural data better than classical models, suggesting that they can capture patterns that are not assumed in cognitive models. However, the black-box nature of large neural networks makes it difficult to extract interpretable cognitive processes from fitted RNNs. JBM addressed this issue by introducing tiny RNNs together with methods to visualise their latent dynamics, thereby proposing a novel framework for analysing learning and choice behaviours.

By contrast, RNNs can flexibly adjust their behavioural dynamics based on recent choices and reward histories, thereby tracking individual differences and within-individual state changes^8^. In RL terms, this involves modulating key parameters such as the learning rate *α*, which controls how strongly action values are updated, and the inverse temperature *β*, which regulates the stochasticity of choice. The ability to adjust predictions based on context, known as in-context learning or in-context adaptation, has been especially valuable in generative AI^9^. In cognitive modelling, however, such flexibility can be problematic, as the objective is to interpret fitted dynamics in relation to underlying cognitive processes^8^.

To illustrate this issue, we simulated a simple reversal learning task, assuming that the agent alternates between engaged and disengaged states, a phenomenon that has been reported in both humans and non-human animals during decision-making tasks^10,11^ (See Supplementary Text 1 for details on simulations). We modelled the agent using a simple model-free RL framework, varying the level of choice randomness (inverse temperature) by state: *β* = 3.0 in the engaged state (less random, more sensitive to value differences) and *β* = 0.2 in the disengaged state (near-random). The resulting ground-truth choice probabilities are shown as grey lines in Figs. 1a and 1b.

**Fig. 1.**
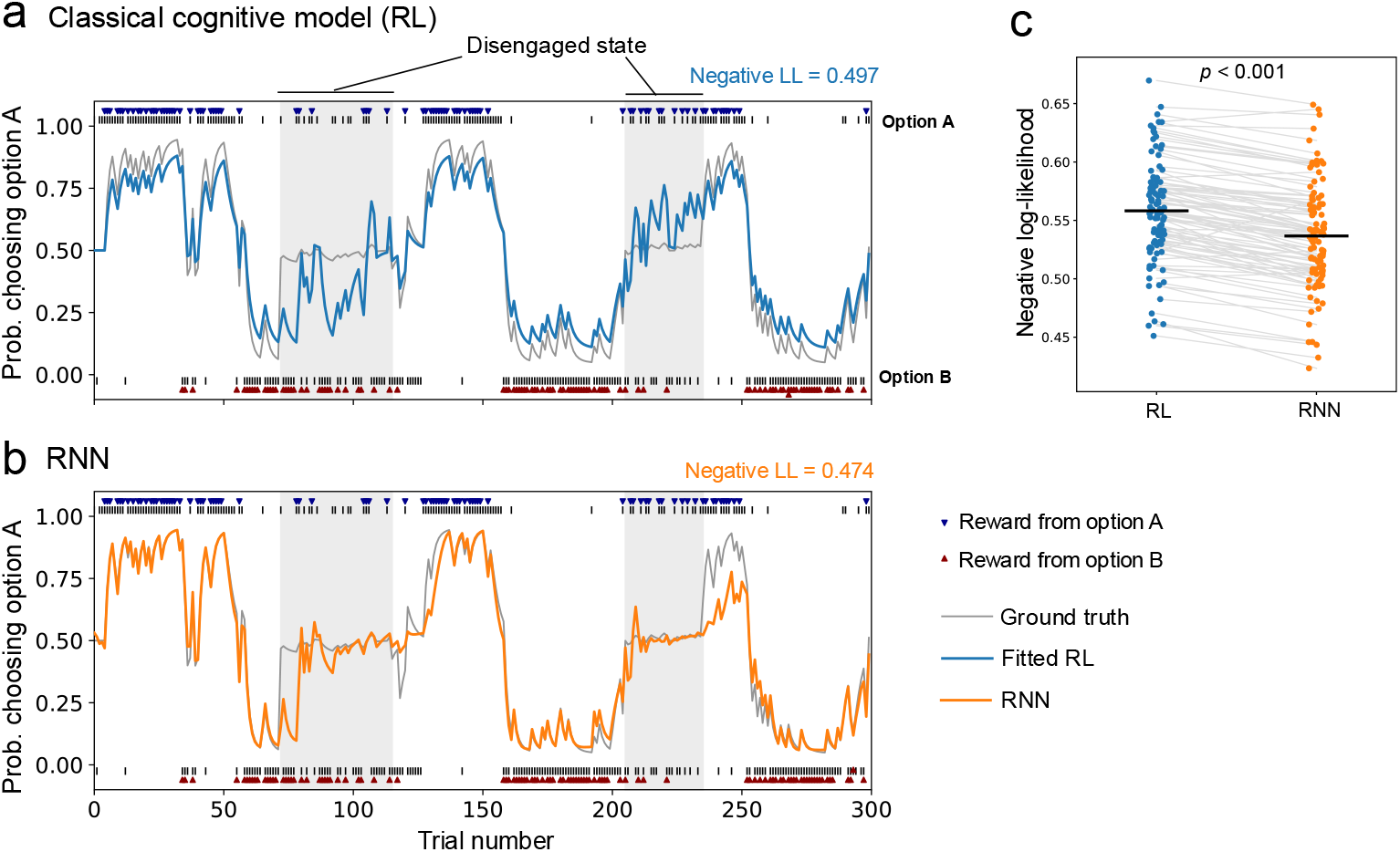
Illustration of how RNNs can track agent state transitions through in-context adaptation. **a, b** Simulated choice data in which the agent alternates between the engaged and disengaged states, along with model predictions. The same data are shown in both panels but separated by a fitted model for clarity. Panel **a** shows predictions from the classical cognitive model (reinforcement learning, RL; light blue line), and panel **b** shows predictions from the recurrent neural network (RNN) with four gated recurrent units (GRUs; orange line). The grey lines indicate the ground truth (choice probabilities from the generative model used in the simulation). These data were obtained from a representative test session that was not used for model fitting. **c** Comparison of negative log-likelihoods (LLs) for RL model and RNN on test data across 100 sessions. Each dot and connecting line represent one session of 300 trials. A paired t-test revealed a significant difference (*t*_99_ = 13.74, *p*< 0.001, *d* = 1.37 [95% CI, 1.10–1.65]), with the RNN showing a higher predictive accuracy. The black horizontal bars indicate the mean across sessions. See Supplementary Text 1 for the details of the simulations.

We then fit these simulated data with a classical RL model and an RNN with four gated recurrent units (GRUs), a configuration that considers a *tiny RNN*. The choice probabilities predicted by an RL model with the same structure as the ground-truth model but using a single constant *β* across all trials are denoted by the blue line in Fig. 1a. Although such a model could fit the data perfectly if the engaged and disengaged states were estimated separately, the difference in *β* across states forces the fitted parameter to take an intermediate value. As a result, in the engaged state the predictions drift toward chance-level choice (≈0.5), while in the disengaged state they drift toward biased choice. By contrast, the RNN predictions (Fig. 1b, orange line) initially deviate from the ground truth after each state switch but quickly converge to the appropriate level (e.g., ≈0.5 in the disengaged state) through in-context adaptation. This flexibility enables the RNN to track state changes and maintain predictions close to the ground truth even in the engaged state. Consequently, the RNN achieves a lower (i.e., better) negative log-likelihood on held-out test data compared with the RL model (Fig. 1c).

RNN achieve superior predictions by modulating the degree of randomness based on recent behaviour, for example, by inferring randomness from recent choice history. However, animals are unlikely to explicitly monitor their past choices in such a manner. A more natural explanation is that internal states governing behavioural properties switch discretely, as modelled using hidden Markov models^10,11^. Therefore, although RNNs can flexibly adapt to their dynamics through in-context adaptation, this property may not directly reflect the cognitive strategies of animals.

The properties of the RNNs described above have important implications for the interpretation of the JBM results. First, although JBM reported that RNNs consistently outperformed classical cognitive models in predictive performance, this advantage may reflect the ability of RNNs to track state changes, which is not available to cognitive models. If this is true, the superior performance of RNNs does not necessarily imply that they capture cognitive processes beyond those represented by cognitive models. Second, and relatedly, the dynamical rules extracted through the JBM’s dynamical systems analyses (e.g. phase portraits and dynamical regression) may also incorporate elements of in-context adaptation to state changes rather than purely cognitive processes.

Unlike animal data, human behavioural data typically include fewer trials per participant, making it difficult to obtain stable estimates when fitting RNNs at the individual level. Classical cognitive modelling addresses this with hierarchical models that assume a population-level distribution and stabilise individual estimates using pooled information^12^; however, such approaches are less feasible for RNNs given their large number of parameters. JBM proposed a “knowledge distillation” framework, which can mitigate this issue. This framework involves training a large teacher network on the entire dataset, incorporating participant identity as an input (i.e. subject-specific embeddings), and then training smaller student networks for individual subjects to mimic the output. These student networks can subsequently be analysed to interpret the underlying dynamics.

Although this approach is rational, it may also be susceptible to the influence of the in-context adaptation of RNNs, namely, their individual-difference tracking (IDT) property. In a teacher network, individual differences are expected to be captured through subject-specific embeddings. However, Song et al. found that such embeddings improved fit only in the first few trials, suggesting that IDT accounted for later differences. If student networks inherit the IDT properties of teachers, their dynamics may not purely reflect cognitive processes, a possibility that warrants further examination.

Rigorous validation is required to establish a novel methodological framework. The framework proposed by JBM is highly promising and could become a general approach in computational cognitive modelling; however, further verification is still needed. The points we raise here are intended to contribute to this process and do not diminish the value of the JBM’s results or the significance of their approach. They highlight issues that should be considered when applying the framework, and addressing them may lead to further refinement.

## Code availability

The Python script used for the simulation presented in Fig. 1 is available at https://osf.io/hnb7q/files/osfstorage/68e21fe2f706d0af1d6d3323.

## Acknowledgements

This work was supported by JSPS KAKENHI Grant Numbers 24K15121 and 25H01173 (K.K.).

## Competing financial interests

The author declares no competing financial interest.

## Supplementary Text

### S1 Simulation setting

The simulation shown in Fig. 1 of the main text largely followed the procedure described in ref. ^1^. The main difference was that the simulated agent was assumed to alternate between engaged and disengaged states. Additionally, motivated by animal experiments where tens of thousands of trials are typically obtained from a single subject, we simulated 200 sessions of 300 trials each for a single agent.

To generate data, we simulated the behaviour of the agent in a reversal learning task (two-armed bandit task). One option was associated with a high reward probability of 0.8 while the other was associated with a low reward probability of 0.2. At each trial *t* (where *t* denotes the trial index), a reward was given (*r*_*t*_ = 1) based on the probability associated with the chosen option; otherwise, no reward was given (*r*_*t*_ = 0). After each 50-trial block, the reward probabilities of the two options were reversed. Each session comprised 300 trials with five reversals, and we simulated 200 sessions in total. Data from 100 sessions were used as training data for fitting the RNN and RL models, and the remaining 100 sessions were used as test data to evaluate the predictive accuracy of these models. All simulations were implemented in Python (version 3.12.1).

### S2 Agent model

The agent was modelled using a forgetting Q-learning (FQ-learning) model ^2,3^. FQ-learning is described by a one-dimensional variable and has been shown to fit behavioural data better than reinforcement learning without forgetting ^2,4–6^. It has also been demonstrated that even a simple RNN with a single linear unit can exactly reproduce the behaviour of FQ-learning in the absence of state changes ^1^, implying that RNNs are expected to learn effectively within each state (engaged or disengaged). Thus, by adopting FQ-learning as the base model of the agent, we highlight the capacity of RNNs to track state transitions ^1^. As shown in Section S3, this model is equivalent to the “model-free strategy (*d* = 1)” used by Ji-An et al. as their basic RL model ^7^.

In the FQ-learning model, the Q-value (action value), *Q*_*t*_(*a*_*t*_), for a chosen option *a*_*t*_ *∈ {A, B}* at trial *t* is updated as

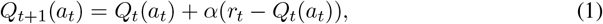

where *α* ∈ [0, 1] is the learning rate, which determines how strongly the prediction error updates the value, and *r*_*t*_ *∈ {*0, 1*}* is the reward received on trial *t*.

The Q-value for the unchosen option *ā*_*t*_ is assumed to decay as

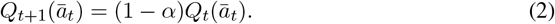

In this model, the learning rate *α* also determines how quickly the value of the unchosen option decays, thereby serving as the forgetting rate.

The choice probability for option A is given by the softmax function:

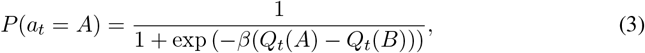

where *β* is the inverse temperature, controlling sensitivity to value differences; larger *β* makes choices more deterministic.

Within each session, the agent alternated between an “engaged state” and a “disengaged state”. State durations were sampled from uniform distributions of 70–100 trials (engaged) and 30–80 trials (disengaged), concatenated until the session reached 300 trials. Each session began in the engaged state. During the engaged state, the inverse temperature was set to *β* = 3.0, yielding relatively deterministic choices favouring the higher-valued option while still allowing occasional exploration. In the disengaged state, *β* = 0.2, producing behaviour close to random choice. The learning rate was fixed at *α* = 0.4, and the initial Q-values for both options were set to zero.

With this setup, the agent necessarily entered the disengaged state at least twice per session. Although this is a simplified assumption, we adopted it to clearly demonstrate that the RNN can track state changes and thereby achieve better fits.

### S3 Fitting RNN and RL

For the RL model, we fit the FQ-learning model with a fixed *β* to the simulated data. In other words, the fitted RL model was assumed to share the same structure as the ground-truth model within each state.

For the RNN, we employed the same architecture described in ref. ^1^. Specifically, the RNN architecture consisted of four gated recurrent units (GRUs ^8^). The input at each trial was a four-dimensional vector representing the action and reward from the previous trial. Actions were one-hot encoded using two input units: the selected action was coded as 1 and the unselected action as 0. Rewards were encoded using a choice-dependent reward (CDR) scheme: for each action, a dedicated input unit was assigned, with the reward value (0 or 1) entered only for the chosen action, while the unchosen action’s input was set to 0. This CDR encoding differs from that used by Ji-An et al. ^7^, where action and reward were each encoded as a single input (together with an additional state input). Moreover, Ji-An et al. also introduced switching RNNs, in which network weights change depending on the input, as one of their candidate models. Our one-hot action encoding and CDR scheme can be regarded as a special case of such switching RNNs, since the connectivity to the GRU is effectively switched depending on the chosen action, while the recurrent connections remain fixed. The output layer consisted of two units, with choice probabilities obtained via a softmax transformation. Further implementation details of the RNN architecture are provided in ref. ^1^.

Both the RNN and RL models were fit using maximum likelihood estimation: for each trial in the 100 training sessions, the model generated the choice probability based on the preceding trial, and parameters were optimised to minimise the negative log-likelihood (i.e., cross-entropy) across all sessions.

For the RNN, it is standard practice to apply early stopping with cross-validation to determine the number of training iterations and avoid overfitting. In this study, however, the true choice probabilities were known from the generative model. We therefore selected the training step at which the RNN’s predicted choice probabilities were closest to the true probabilities. Specifically, across 20,000 training steps, we computed the Kullback–Leibler (KL) divergence between predicted and true probabilities every 100 steps and adopted the model from the step at which this divergence was minimized (13,600 steps). This approach eliminated dependence on a particular validation dataset and allowed us to capture the RNN’s optimal behaviour. In contrast, the RL model had only two free parameters, making overfitting unlikely; no such procedure was applied.

The negative log-likelihood for the test data was then calculated to compare predictive accuracy between models.

### S4 Equivalence between FQ-learning and the one-dimensional model-free RL

Here, we show that the FQ-learning model is equivalent to the basic RL model used by Ji-An et al. ^7^, namely the model-free strategy (*d* = 1).

According to Eq. (3), the choice probability depends only on the difference between the Q-values. Thus, to characterize the model’s behaviour, it is sufficient to describe the dynamics of this difference ^3^. Let this difference be defined as *h*_*t*_ = *Q*_*t*_(*A*) *− Q*_*t*_(*B*).

Here, Eq. (1) and Eq. (2) can be written as

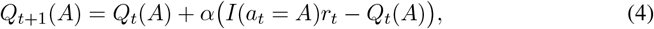

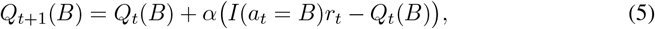

where *I*(·) denotes the indicator function, which takes the value 1 if the statement inside is true and 0 otherwise.

Subtracting Eq. (2) from Eq. (1) gives the update rule for *h*_*t*_:

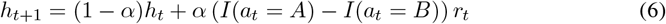

Thus, the dynamics can be expressed as a one-dimensional variable ^3^.

The model-free strategy (*d* = 1) of Ji-An et al. ^7^ can be written, after a slight rearrangement,as

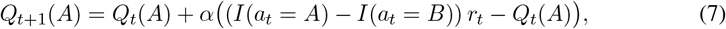

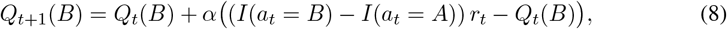

By subtracting the second equation from the first, we obtain

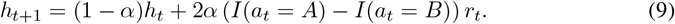

Compared with Eq. (6) for FQ-learning, the only difference is a factor of 2. Since the Q-values are initialized uniformly (in our simulation, *Q*_1_(*A*) = *Q*_2_(*B*) = 0), the initial value of *h*_1_ is zero. Taking both the factor of 2 and the initial conditions into account, this difference can be absorbed by halving *β*, rendering the two models equivalent.

## Notes

### Competing Interest Statement

The authors have declared no competing interest.

